# Mammalian Esophageal Stratified Tissue Homeostasis is Maintained Distinctively by the Epithelial Pluripotent p63^+^Sox2^+^ and p63^−^Sox2^+^ Cell Populations

**DOI:** 10.1101/2023.03.22.533592

**Authors:** Xiaohong Yu, Hui Yuan, Yanan Yang, Wei Zheng, Xuejing Zheng, Shih-Hsin Lu, Wei Jiang, Xiying Yu

## Abstract

Self-renewing, damage-repair and differentiation of mammalian stratified squamous epithelia are subject to tissue homeostasis, but the regulation mechanisms remain elusive. Here, we investigate the esophageal squamous epithelial tissue homeostasis in vitro and in vivo. We establish a rat esophageal organoid (rEO) in vitro system and show that the landscapes of rEO formation, development and maturation trajectories can mimic those of rat esophageal epithelia in vivo. Single-cell RNA sequencing (scRNA-seq), snap-shot immunostaining and functional analyses of stratified “matured” rEOs define that the epithelial pluripotent stem-cell determinants, p63 and Sox2, play crucial but distinctive roles for regulating mammalian esophageal tissue homeostasis. We identify two cell populations, p63^+^Sox2^+^ and p63^−^Sox2^+^, of which the p63^+^Sox2^+^ population presented at the basal layer is the cells of origin required for esophageal epithelial stemness maintenance and proliferation whereas the p63^−^Sox2^+^ population presented at the suprabasal layers is the cells of origin having a dual role for esophageal epithelial differentiation (differentiation-prone fate) and rapid tissue damage-repair responses (proliferation-prone fate). Given the fact that p63 and Sox2 are developmental lineage oncogenes and commonly overexpressed in ESCC tissues, p63^−^Sox2^+^ population could not be detected in organoids formed by esophageal squamous cell carcinoma (ESCC) cell lines. Taken together, these findings reveal that the tissue homeostasis is maintained distinctively by p63 and/or Sox2 dependent cell lineage populations required for the tissue renewing, damage-repair and protection of carcinogenesis in mammalian esophagi.

## Introduction

The mammalian esophagus contains a stratified squamous epithelium that is originated from the dorsal side of the anterior foregut during embryotic development. The anterior foregut first forms a simple columnar epithelium and then specifies the esophagus off from the respiratory system by a process called respiratory-esophageal separation (RES) [1]. After RES and embryotic development proceeded, the esophageal epithelium is transformed into a multilayered stratified squamous epithelium including the basal layer(s), the suprabasal layers and differentiated/keratinized layers [2]. The multilayered stratified squamous epithelium is persistent throughout adulthood that functions as a barrier from the lumen [3]. Hence, it is essential for the esophageal epithelium kept in an exquisite balance of proliferation-differentiation homeostasis, damage-repair response, and disease protection as the epithelium of the adult esophagus turns over twice a week in rodents and once in two weeks in humans [4].

The underlying mechanisms by which the esophageal specification, RES and esophageal epithelial morphogenesis during the development are regulated and determined have been extensively investigated in various model systems in vivo and in vitro [5, 6]. Accumulating evidence indicates that multiple signaling pathways and factors, especially the stemness pluripotent transcription factors, p63 and Sox2, play critical roles in regulating and controlling these processes [1]. Sox2, one of sex-determining region Y [SRY]-box family transcription factors, is essential for foregut stem cell pluripotency [7, 8]. High levels of Sox2 stimulate the formation of stratified squamous epithelium and prevent pseudostratified columnar epithelium, lining the trachea [9]. Thus, high expression of Sox2 is detected in the proliferative basal layer of the stratified squamous epithelium [5, 7, 8, 10]. Consistently, dysregulated overexpression of Sox2 functions as a lineage-survival oncogene in human cancers including esophageal squamous cell carcinomas (ESCCs) [11, 12]. Gene amplification and overexpression of Sox2 is closely associated with poor prognosis in patients with ESCCs and other squamous cell carcinomas (SCCs) [13]. The subtype of p53-related p63, ΔNp63 (p63), is determined as a crucial “switch” required for the cell lineage specifications and tissue proliferation-differentiation homeostasis during the esophageal development [14, 15]. P63 functions as a potent regulator whose expression results in the proliferation of the basal layer and the stratification of the epithelium [16]. Notably, human p63 gene is located at chromosome 3q28, near to Sox2 gene locus, which is often co-amplified and co-overexpressed with Sox2 gene in ESCCs and other SCCs. Thus, Sox2 and p63 are thought to be the important cooperative partners involved in controlling the esophageal stratified squamous epithelial homeostasis and carcinogenesis, but the precise mechanisms remain elusive [11, 12, 17].

It is much less understood how the multilayered stratified (“matured”) esophageal squamous epithelium in an adult maintains its tissue homeostasis and what the underlying mechanisms of tissue homeostasis maintenance, damage/stress/injury-repair/recovery and disease-related perturbations including esophagitis and ESCC are controlled, regulated and/or developed. Given the fact that the stratified esophageal epithelium is often attacked by chemical/mechanical stresses and acute/chronic inflammatory conditions during life, ultimately leading to irreversible damages and illnesses, many laboratories including us have made a great deal of efforts by establishing/utilizing in vitro and in vivo model systems, e.g. cell and organoid cultures from esophageal keratinocytes, transgenic mice, carcinogen-induced esophageal carcinogenesis in rodents and human samples from esophagitis and ESCC patients to seek for solutions [4, 18–23]. Recently, we and others demonstrate that the mammalian matured esophageal epithelia display the heterogeneity at the single-cell level [1, 24–26]. A small population of cells with high levels of hemidesmosomes (HDs) has stemness potential in the context of tissue homeostasis and aging in the basal layer of stratified epithelium [24, 25, 27]. However, the cells of origin that control proliferations, differentiations and damage-repair responses in esophageal stratified matured squamous epithelia have not been clearly determined.

The in vitro three-dimensional (3D) organoid culture platforms have emerged as powerful experimental tools and/or systems for investigating tissue and organ development, morphogenesis, structural maintenance, proliferation-differentiation homeostasis and disease-related perturbations including cancer [28]. Currently, mammalian esophageal organoids (mEOs) can be derived successfully from pluripotent stem cells (PSCs) or esophageal progenitor/stem keratinocytes from the basal layer, which spontaneously organize into multilayered stratified squamous epithelia. The stratified epithelial structures of mEOs enriched with cell types and tissue layers that typically present in the esophageal epithelial tissue of origin in vivo [9, 18, 29–33]. Hence, mEOs have been utilized to investigate esophageal tissues homeostasis maintenance and disease-related problems in vitro. Suppression of the Notch signaling-mediated squamous cell differentiation by cytokines could mimic the phenotypic alterations in eosinophilic esophagitis (EoE), a chronic inflammation disease in the human esophagus [32, 34, 35]. In vitro mEOs could be particularly useful for determining how a multilayered stratified squamous tissue formation, self-renewal, proliferation-differentiation homeostasis, damage/stress/injury-repair/recovery, and how a disease-related esophagitis and ESCCs are regulated, which could not be easily done in vivo [20, 33, 36–38].

In this study, we report to establish a stable in vitro rat esophageal organoid (rEO) model system and use rEO system together with rat animals to investigate the esophageal squamous stratified epithelial tissue homeostasis. We demonstrate that esophageal tissue homeostasis is maintained by two epithelial pluripotent cell populations, p63^+^Sox2^+^ and p63^−^Sox2^+^, respectively.

## Materials and methods

### Establishment of immortalized normal rat esophageal keratinocyte cell line

Human telomere reverse transcriptase (hTERT) immortalized rat normal esophageal epithelial cell line (RNE-D3, D3 for short) were established and preserved by our laboratory. Primary esophageal basal keratinocytes from males of Rat F344 strain were isolated and then infected with retrovirus made by pBABE-hTERT. After the puromycin selection, several rat esophageal keratinocyte cell lines expressing hTERT including the D3 cell line were cloned and expanded in vitro. D3 cells were cultured in DMEM/F12 (3:1) medium supplemented with 10% fetal bovine serum (Thermo Fisher Scientific), 8 ng/mL Cholera Toxin (CELL technologies), 5 ng/mL insulin (CELL technologies), 25 ng/mL hydrocortisone (CELL technologies), 0.1 ng/mL EGF (PeproTech) and 10 μM Y27632 (Topscience) in a humidified 37 °C incubator supplemented with 5% CO2.

D3-shSox2 cells were constructed by lentivirus transduction using following sequences: shRNA-1: 5’-CCAAGACGCUCAUGAAGAAGG-3’. shRNA-2: 5’-GCUACAGCAUGAUGCAGGAGC-3’. shRNA-3: 5’-GGGACAUGAUCAGCAUGUACC-3’. D3-shp63 cells were constructed by lentivirus transduction using following sequences: shRNA-1: 5’-ACUGUAGGGUAGCACACCGUG-3’. shRNA-2: 5’-UCAAUCUUGUUUGUUGCACCA-3’. shRNA-3: 5’-UGGAAGGACACAUCGAAGCUG-3’ (Beijing Syngentech Co., LTD).

D3-Sox2 overexpression cells were constructed by lentivirus transduction using following plasmid: pLV-Neo-SOX2-HA (PolePolar Biotechnology Co., LTD).

### D3-organoids culture, passage, freezing, and thawing

Single cells isolated from D3 cell line were counted and then a total number of 1000 cells were mixed with 50 μL of Matrigel (Corning) and plated in 24-well plates. After polymerization of the Matrigel, advanced DMEM/F12 containing R-spondin1 (R&D Systems), Noggin (Pepro Tech) and ROCK inhibitor Y-27632 (Sigma Aldrich) was added and changed every 2 days. For passage, D3-organoids were removed from the Matrigel, which dissolved by adding cell recovery solution, and then transferred to fresh Matrigel after proteolysis of individual cells. Passage was performed every week with a 1:3-1:5 split ratio. Organoids were digested into single cells with TryPLE (Thermo Fisher Scientific) and dispersion in 90% FBS with 10% DMSO (Sigma Aldrich) in −80°C. The size of organoids was measured by ImageJ software from representative images.

### Rat ESCC Cell lines (RESCs) and rat ESCC organoids culture

Rat esophageal squamous carcinoma cell lines (RESCs) were driven from the an NMBzA-induced rat ESCC model of our lab. Several RESCs were cloned and expanded. RESC-1 was selected and cultured in RIPM160 medium supplemented with 10% fetal bovine serum (FBS).

For 3D organoids culture, RESC-1 cells were trypsinized to prepare cell suspensions. Single cells were counted and a total number of 2000 cells were mixed with 50 μL of Matrigel (Corning) and plated in 24-well plates. After solidification, 500 μL of advanced DMEM/F12 (Thermo Fisher Scientific) supplemented with 1 × N2 (Thermo Fisher Scientific), 1 × B27 supplements (Thermo Fisher Scientific), 0.1 mM N-acetyl-L-cysteine (Sigma-Aldrich), 50 ng/mL recombinant human EGF (R&D Systems), 2% Noggin/R-Spondin–conditioned media (R&D Systems), 100 ng/mL recombinant human Wnt3A (R&D Systems), 500 nM A83-01 (R&D Systems), 10 nM SB202190 (R&D Systems), 10 nM gastrin (R&D Systems), 10 nM nicotinamide (Sigma-Aldrich), and 10 μM Y27632 (Selleck), was added and replenished every other day. RESC-1 organoids were grown for 13-16 days for analysis.

### Organoid formation rate and organoid size analysis

After 10 days of organoid formation by primary esophageal cells or immortalized D3 cell line in Matrigel (Corning), the number and size of organoids (spheres larger than 50 μm in diameter) were quantified using Image J software. Organoid forming efficiency was defined as: the number of organoids/number of cells seeded in Matrigel multiply by 100%. The size of each organoid was analyzed by measuring the diameter under the microscope. All experiments calculated at least three parallel groups.

### H&E and immunofluorescence analyses

H&E and immunofluorescence analyses were performed on OCT-embedded from primary esophageal epithelial tissues or organoids as previously descried [24]. Briefly, the OCT compound-embedded esophageal tissue or organoids were cut into 5-10 µm sections using a cryosection system. Mayer’s hematoxylin and 1% eosin Y solution (Servicebio) were used for H&E staining.

For immunofluorescence analysis, antigens were retrieved using EDTA antigen retrieval solution, and then blocked with 10% normal goat serum (containing 0.2% TritonX-100) for 1 hour at room temperature. Primary antibodies against Sox2 (Abcam, 1:500), p63 (Abcam, 1:100), Cytokeratin14 (Abcam, 1:1000), Cytokeratin13 (SantaCruz, 1:500) were used to incubate overnight. The slides were washed three times and incubated with corresponding secondary antibodies (goat anti-rabbit secondary antibodies Alexa Fluor 488, or goat anti-mouse secondary antibodies Alexa Fluor 568, Invitrogen, 1:300) for 1 hour at room temperature. Samples were counterstained with DAPI and then washed three times with PBS, and the slides were mounted with prolong gold anti-fade mounting medium (Invitrogen), sections were imaged using a fluorescence microscope (Leica). All the images were converted to 8 bit grayscale for plot profile analysis.

### Establishment of rat esophageal acute inflammatory model

F344 rats, 7-10 weeks old, were purchased from BEIJING HUAFUKANG BIOSCIENCE COMPANY. Briefly, Rats were treated daily with 1 nmol MC903 (Tocris Bioscience) on ears in the presence of 50 ug OVA (Sigma-Aldrich) for 14 days. Rats were challenged intragastrically with 25 mg OVA on day 15 and 18. Upon the first intragastric OVA challenge, rat were continuously provided with water containing 0.75g/L OVA. All rats were euthanized on day 20. The entire esophagi were obtained and fixed by 10% formalin overnight, followed by routine histological processing of H&E. For immunofluorescence, the entire esophagi were fixed by 10% formalin overnight and dehydrated in 30% sucrose solution. Then the segmented esophagi were embedded in OCT compound and immediately frozen by liquid nitrogen to proceed with cryosection.

### Generation of inflammatory damage/stress-repair esophageal organoids by inflammatory factor TGF-α and IL-13

D3 single cells were suspended and cultured in Matrigel (Corning) for 5 days or 7 days before treatment with TGF-α (R&D, 40 ng/mL) or IL-13 (R&D, 20 ng/mL) for 5 days or 3 days. To allow the acute inflammatory stress response recovery, D3-organiods were cultured to Day 5 and then treated with TNF-α (R&D, 40 ng/mL) and IL-13 (R&D, 20 ng/mL) for 3 days to induce acute inflammatory stress responses. At Day 8, D3-organiods were withdrawn the TNF-α and IL-13 treatment and replaced into fresh medium for 2 more days.

### Generation of shSox2/shp63 (3.5) rat esophageal organoids

After 3.5 days of organoid cultured by immortalized D3 cell line in Matrigel (Corning), lentivirus shSox2 or shp63 were transient transfected with MOI of 100 in 3D culture system for 2 days. The organoids were changed to fresh organoid culture medium, which were containing advanced DMEM/F12 with R-spondin1 (R&D Systems), Noggin (Pepro Tech) and ROCK inhibitor Y-27632 (Sigma Aldrich). Then the shSox2/shp63 (3.5) esophageal organoids were grown for more 4.5 days for analysis.

### Protein extraction and Western blotting analysis

Cells or organoids of 10-20 mg was homogenized in 200 μL RIPA supplemented with protease and phosphate inhibitors (ThermoFisher) at 4°C. Cell lysate samples were centrifuged at 15000 RPM for 15 minutes at 4°C, and protein concentration was measured using a BCA protein assay kit (ThermoFisher). Equal amount (30 μg per lane) of total denatured proteins was loaded onto a Criterion 12-well gel and subjected to immunoblotting. The primary antibodies and dilutions were used against Sox2 (Abcam, 1:1000), p63 (Abcam, 1:1000), S100a8 (beyotime,1:1000), S100a9 (beyotime, 1:1000), Cyclin D1 (Abcam, 1:1000), ECT2 (Abcam, 1:1000), KLF4 (Abcam, 1:1000), INC1 (CST, 1:1000), CK13 (Abcam, 1:1000), CK14 (Abcam, 1:1000), Bax (Beyotime, 1:1000), Cleaved-caspase3 (Beyotime, 1:1000), Cleaved-PARP (Beyotime, 1:1000), NF-kB (CST, 1:1000) and β-actin (beyotime,1:1000), respectively, and incubated for overnight, followed by developing with secondary antibodies (Beyotime, 1:5000) for 1 hour at room temperature. Immunoblots were visualized with Cham Chemi (Sagecreation).

### Statistical analysis

Student’s t-test and Two-way ANOVA were performed for analyzing statistic differences between groups, and p<0.05 was consider significance. Data were presented as Mean±SD. GraphPad was used for analysis.

## Results

### Establishment of a stable in vitro rat esophageal organoid (rEO) model system

We aimed to establish a stable in vitro organoid model system that could investigate the maintenance of esophageal tissue homeostasis in detail. To achieve our goal, we first isolated primary keratinocytes from the basal layers of rat esophagi and then cultured them either directly into organoids in Matrigel (3-D) or briefly plated these cells in petri dishes (2-D) followed by organoid cultures. Under the organoid culture condition we modified, the isolated primary rat esophageal basal keratinocytes could form organoids with stratified squamous layers: 1) the basal layer at the outermost, 2) the suprabasal layers in the middle and 3) the keratinization layer at the center, in 10-12 days (Supp. Fig. 2a). However, these primary esophageal keratinocytes could not be continually maintained in the 2-D and 3-D culture conditions after 3-5 passages.

To overcome the problem, we then decided to generate human telomerase reverse transcriptase (hTERT) immortalized normal rat esophageal keratinocyte cell lines using retrovirus mediated transduction (for detail, see experimental procedures) (Fig. 1a). To this end, several clonal rat normal esophageal epithelial cell lines (RNEs) expressing hTERT were cloned and expanded in vitro. We focused on three cell lines, REN-A1, REN-D3 and REN-F3 (Supp. Fig. 1a). These cell lines could grow continually in the 2-D culture condition over 100 passages in vitro but did not form colonies in soft agar and tumors in nude mice (Fig. 1b). Immunofluorescence demonstrated that, under the 2-D cell culture condition, these hTERT-immortalized cell lines expressed the esophageal basal cell marker CK14 but not the suprabasal cell marker CK13 (Fig. 1c and Supp. Fig. 1b). After long-term cultures (passages >50), the morphologies of these cell lines remained the same, like those of their early passages (<10 passages) with typical cobblestone shapes (Fig. 1d). Notably, when compared with the rat esophageal primary epithelial keratinocytes from the basal layer, not only early passages but also late passages of these immortalized cell lines did not alter the gene expressions of several key cell regulatory pathways that controlled cell proliferation-differentiation (H-ras, CK13, CK14, Notch1, VIM, CDH2 and FN1), cell cycle progression (Cyclin D1, CDK4/6, p-RB, p21 and MCM3), stemness maintenance (Sox2, p63 and β-catenin) and stress response and apoptosis (p53, CHK1, CHK2 and γ-H2AX) determined by immunoblotting (Fig. 1e, f), immunofluorescence (Fig. 1c and Supp. Fig. 1b, d) and gene cluster (Supp. Fig. 1e, f) analyses. We also performed karyotype and whole genomic sequencing analyses using REN-D3 (D3 for short) cell line. The results showed that, beside an integrated hTERT sequence in rat chromosome 17q, D3 cells had a normal rat genome with 42 chromosomes, demonstrating that D3 was an hTERT-immortalized rat “normal” esophageal squamous keratinocyte cell line (Supp. Fig. 1c, g).

**Fig 1.**
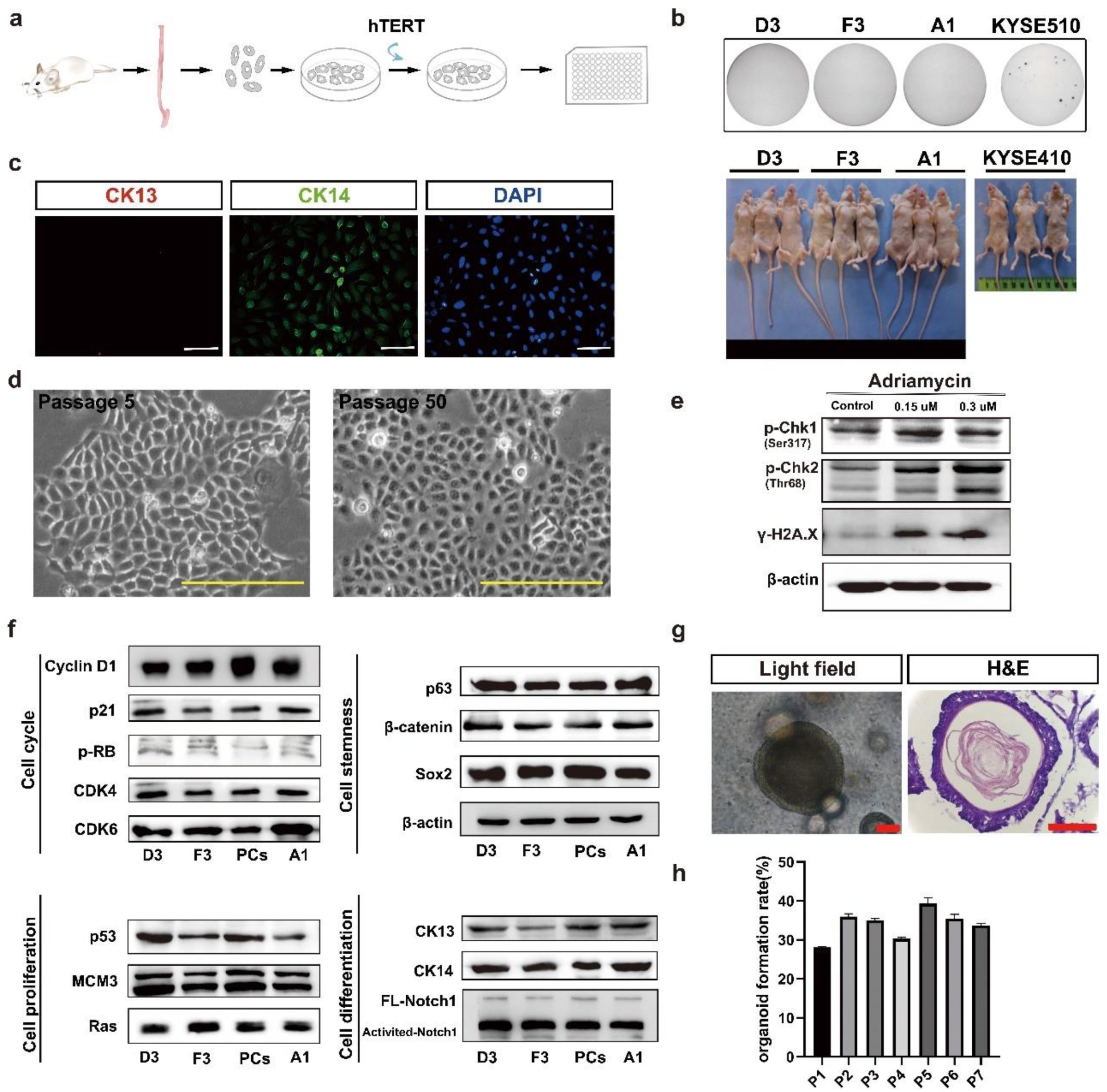
Establishment of a rat esophageal organoid (rEO) model system in vitro. **a** Schematic illustration of the establishment of rat normal immortalized esophageal epithelial cell lines (RNEs) in 2-D. **b** Representative images of soft agar-stained and tumor formation in nude mice of RNE cell lines D3, F3 and A1, with human ESCC cell lines KYSE410 and KYSE510 as positive controls. **c** Immunofluorescence staining of CK13 and CK14 in D3 cell line, Scale bar: 100 μm. **d** Representative brightfield of D3 cell line at passages 5 and 50, Scale bar: 100 μm. **e** Western blotting of DNA damage repair pathway of D3 cell lines treatment with 0.15 μM and 0.3 μM adriamycin after 12 hours. **f** Western blotting in cell proliferation, cell differentiation, cell stemness, and DNA damage repair pathways among RNE cell lines D3, F3, A1 and primary rat esophageal squamous keratinocyte cells (PCs). **g** Representative brightfield and HE staining of esophageal organoids derived from D3 cell line at Day 10, Scale bar: 200 μm. **h** The organoid formation rate (OFR) of esophageal organoids derived from the D3 cell line at passage 1-7.

We determined whether D3 cells could form stable rEOs in vitro. The organoid formation of D3 cells under our 3-D culture condition was examined, showing that, similar to rat esophageal primary epithelial keratinocytes, D3 cells could grow in Matrigel forming organoids in 10-12 days (Fig. 1g). D3-organoids also displayed the typical stratified squamous epithelial structures with organoid formation rate (OFR) around ∼25% (Fig. 1h). Notably, D3-organoid formation was independent to the culture time and passages of D3 cells in their 2-D culture conditions. Moreover, immunofluorescence analyses of p63, Sox2, CK14 and CK13 epithelial cell markers (see below) demonstrated that D3-organoids had similar staining patterns of rat esophageal stratified squamous epithelia and primary rat esophageal basal keratinocytes derived organoids (Supp. Fig. 2). Taken together, these results demonstrated that establishment of a stable in vitro rEO model system could be achieved using D3 cell line.

### The landscape of D3-organoid formation, development and maturation trajectory in vitro

We examined the in vitro D3-organoid system in detail. Since time-lapse microscopy used for observing long-term organoid cultures (∼10 days) was not applicable even with the light-sheet microscopy, we decided to conduct daily spatiotemporal snapshot imaging plus with statistical analyses of D3-organoids from day 1 to day 10 by various microscopic methods to depict the landscape of D3-organoid formation, development and maturation trajectory in vitro. First, phase-contrast microscopy was applied to snapshot single D3 cells trypsinized from 2D cultures and grown in Matrigel every day for 10 days. As shown in Figure 2A, single D3 cells that in Day 1, first formed small cell clumps in Day 2 to Day 4, which we categorized this stage as organoid formation; then, the cell clumps grew into smoothed and multilayered cell spheres in Day 5 to Day 8, which we categorized this stage as organoid development; and finally, the cell spheres progressed into mature organoids that had tissue-like structures at the out-layers and the keratinized structures in the centers, which we categorized this stage as organoid maturation. Then, we analyzed hematoxylin-eosin (H&E)-stained frozen sections of D3-organoids obtained from Day 2 to Day 10. Consistent with the observations by phase-contrast microscopy, H&E-stained frozen sections of D3-organoids also revealed that D3 organoid formation, development and maturation processes at cellular levels (Fig. 2a) were similar to the esophageal epithelium formation, development and maturation in vivo in rodents reported previously [29]. Notably, under our organoid culture condition, single D3 cells formed small clumps consisting of 6-7 cells with obvious asymmetrical orientations at Day 2 to Day 3. At Day 4 to Day 5, D3-organoids grew to form simple or pseudo-simple columnar epithelia without any sights of cell keratinization. In contrast, at Day 6 to Day 8, D3 organoids developed into smoothed and multilayered cell spheres with a structure in the centers of spheres beginning keratinization and cells at the out-layers organizing into multilayered (1 to 4 layers) stratified squamous tissue-like structures. At Day 9 to Day 10, D3-organoids further matured into multilayered stratified squamous tissue-like structures with a single basal layer-like structure in the outermost-layer (termed as +1 layer), the suprabasal layer-like structures in the inner-layers (termed as +2 to +4 layers) and the keratinized layer in the center of organoids (Fig. 2a, e).

**Fig 2.**
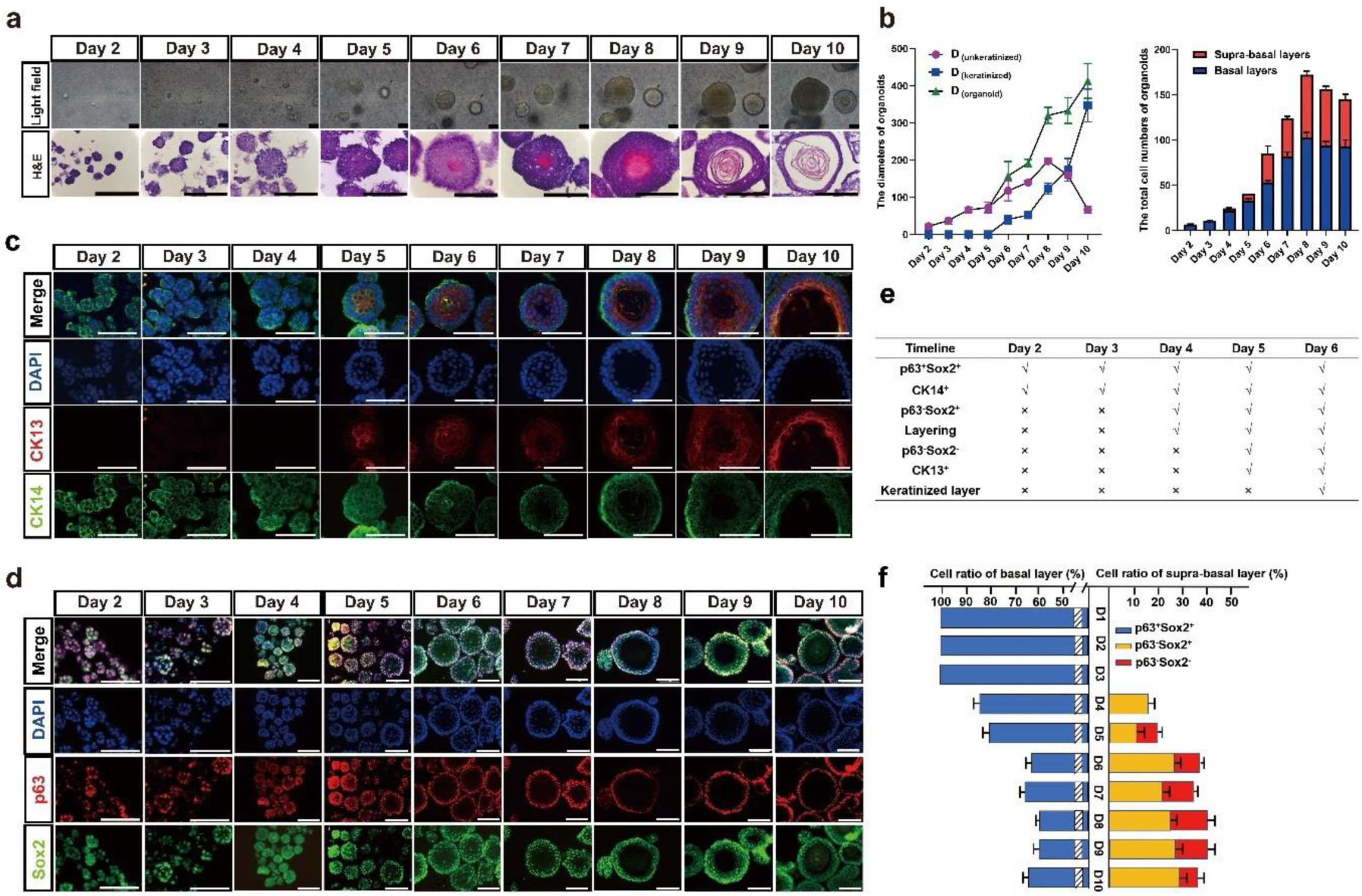
The landscape and dynamics of esophageal stratified epithelial formation trajectory. **a** Representative brightfield and HE staining of esophageal organoids derived from D3 cell line of formation trajectory from Day 1 to 10, Scale bar: 100 μm, 200 μm. **b** The diameters and total cell number of D3-organoids of formation trajectory (n=4, each n represents four random microscope fields, 400×). The data are presented as mean ± SD for percentage analysis, \**P*< 0.05, \*\**P* < 0.01, \*\*\**P*< 0.001, one-way ANOVA test on all pairwise combinations. **c** CK13 (red) and CK14 (green) coimmunostaining of D3-organoids counterstained with DAPI (blue) in formation trajectory, Scale bar: 200 μm. All images are representative of three experiments. **d** P63 (red) and SOX2 (green) coimmunostaining of D3-organoids counterstained with DAPI (blue) in formation trajectory, Scale bar: 200 μm, 400 μm. All images are representative of three experiments. **e** The chart of features of rEO model by formation trajectory. **f** The cell ratio of p63^+^Sox2^+^, p63^−^Sox2^+^ and p63^−^Sox2^−^ populations in the trajectory of esophageal organoids (n=4, each n represents four random microscope fields, 400×). The data are presented as mean ± SD for percentage analysis.

Consistently, the diameters (sizes) of D3-organoids from day 2 to day 10 increased significantly, especially the diameters of the keratinized layers (the transparent centers) of D3-organoids increased dramatically from day 6 to day 10 (Fig. 2b, e). Analysis of H&E-stained organoid cross-sections revealed that the unkeratinized tissue structures (the tissue layers with nucleated cells) of D3-organoids were thickened only from day 2 to day 8 and then became thinned from day 9 to day 10 (Fig. 2b). Numbers of nucleated cells in D3-organoids were 6-7 cells/ORC (organoid cross-section) at Day 2 and then increased to 180 ± 7.6 cells/ORC at Day 8, reaching to maximum. In contrast, numbers of cells in D3-organoids decreased to 165 ± 7.9 cells/ORC at Day 9 and 131 ± 4.4 cells/ORC at Day 10 (Fig. 2b). These results indicated that cell proliferation and differentiation during D3-organoid formation, development and maturation were asynchronized, in which cells only proliferated during organoid formation; cells proliferated and differentiated but the proliferation rates were greater than the differentiation rates during organoid development; and then cells proliferated and differentiated but their proliferation and differentiation rates finally reached to homeostasis during organoid maturation.

To better understand the dynamics of tissue homeostasis, especially proliferation-differentiation at cellular levels, during formation, development and maturation of D3-organoids, we next performed immunofluorescence analyses using various squamous epithelial cell markers, including the endoderm undifferentiated keratinocyte marker cytokeratin14 (CK14), the endoderm differentiated keratinocyte marker cytokeratin13 (CK13), the stemness pluripotent markers p63 and Sox2, and DNA marker DAPI [24]. Consistent with the results obtained from the daily snapshots of phase-contrast microscopy and H&E-stained organoid cross-section analyses, immunofluorescence results revealed that D3-organoids at Day 2 to Day 3 grew as asymmetric small clumps, in which all cells were stained positively for p63, Sox2 and CK14 but negatively for CK13 (Fig. 2c-f and Supp. Fig. 3d). In contrast, D3-organoids began to stratify and formed 1 to 2 epithelial-like layer(s) at Day 4. The p63^+^Sox2^+^CK14^+^ cells were detected in all cells at the outermost layer (+1 layer). However, a few of p63^−^Sox2^+^CK14^+^ cells could be detected at the just stratified +2 layer (Fig. 2c-f and Supp. Fig. 3d). At Day 5, as cells at the stratified +2 layer became CK13^+^ and CK14^−^ cells, the p63^−^Sox2^+^ cells were expressed CK13 instead of CK14 (Fig. 2c-f and Supp. Fig. 3d). Furthermore, a comparison of immunostaining of D3-organoids from day 4 and day 5 revealed that p63^−^Sox2^+^CK13^+^ cells were detected earlier than p63^−^Sox2^−^CK13^+^ cells at the stratified +2 layer, indicating that p63^−^Sox2^+^ cells gradually differentiated into p63^−^Sox2^−^ cells (Fig. 2c-f and Supp. Fig. 3d). Similar results were also observed in D3-organoids at Day 6 to Day 10 although D3-organoids at these time-points growing into unkeratinized stratified multilayered squamous epithelia at the outsides of the spheres and the keratinized layer at the center of the spheres (Fig. 2c-f and Supp. Fig. 3d). Taken together, these results indicated that p63^+^Sox2^+^CK14^+^ cells represented proliferating cells of origin whereas p63^−^Sox2^+^CK14^+^ cells represented differentiating cells of origin in stratified epithelia of D3-organoids. Hence, the “cell fate” and/or “cell lineage trajectory” of D3-organoid formation, development and maturation were single D3 cells proliferating to p63^+^Sox2^+^CK14^+^ cells; p63^+^Sox2^+^CK14^+^ cells proliferating/differentiating to p63^−^Sox2^+^CK14^+^, then to p63^−^Sox2^+^CK13^+^ cells; and finally to p63^−^Sox2^−^ CK13^+^ cells and the keratinized layer, depicting the landscape of stratified epithelial cell proliferation-differentiation and tissue homeostasis of D3-organoids in vitro (Fig. 3a and Supp. Fig. 3e).

**Fig 3.**
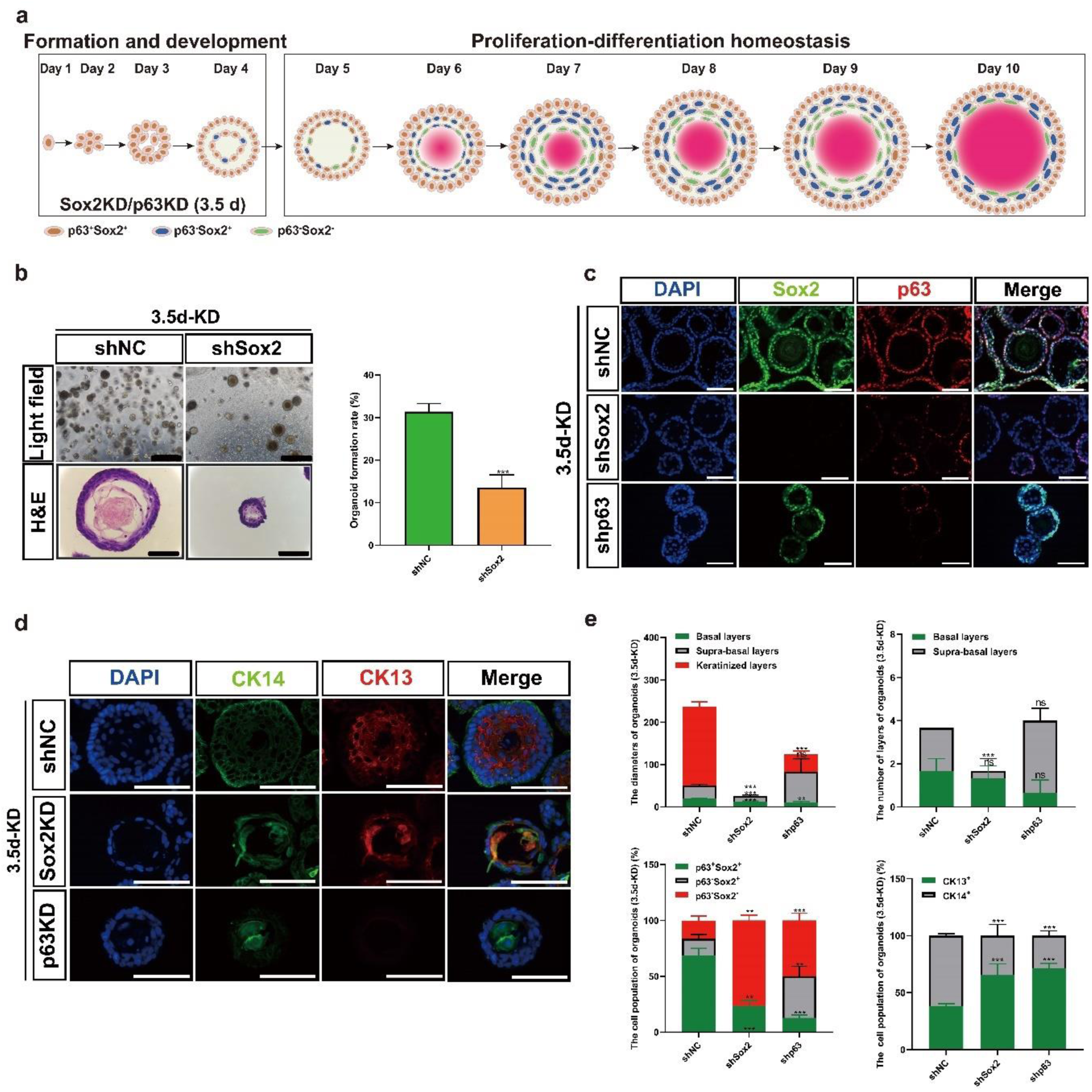
Characterization of the p63^+^Sox2^+^ and p63^−^Sox2^+^ cell populations involved in tissue homeostasis in stratified matured D3-organoids. **a** Schematic illustration of rEO formation trajectory. Sox2 or p63 knockdown by lentivirus in rEO at Day 3.5. **b** Representative brightfield, H&E staining images and quantification of OFR of esophageal organoids at Day 10, Scale bar:200 μm. **c** P63 (red) and Sox2 (green) coimmunostaining of D3-organoids counterstained with DAPI (blue) after Sox2 or p63 knockdown at Day 3.5, Scale bar: 200 μm. All images are representative of three experiments. **d** CK13 (red) and CK14 (green) coimmunostaining of D3-organoids counterstained with DAPI (blue) after Sox2 or p63 knockdown at Day 3.5, Scale bar: 200 μm. **e** Quantifications of the diameters, layers number, cell number of p63^+^Sox2^+^, p63^−^Sox2^+^ and p63^−^Sox2^−^ populations, and cell number of CK13^+^ and CK14^+^ populations of D3-organoids (n=4, each n represents four random microscope fields, 400×). The data are presented as mean ± SD for percentage analysis, \**P*< 0.05, \*\**P* < 0.01, \*\*\**P*< 0.001, one-way ANOVA test on all pairwise combinations.

### Characterization of the p63^+^Sox2^+^ and p63^−^Sox2^+^ cell populations involved in tissue homeostasis in stratified “matured” D3-organoids

Determination of the landscape of D3-organoid formation, development and maturation trajectory enabled us to focus on applying this D3 rEO model system to investigate cell proliferation-differentiation and tissue homeostasis in stratified matured esophageal epithelia, which had not been extensively studied due to lack of an available system. First, we reevaluated the single-cell RNA sequencing (scRNA-seq) data of D3-organoids at Day 10 [24]. We identified that, unlike previously reported that p63^+^ cells were Sox2^+^ cells and vice versa in esophageal epithelia [9]. Sox2^+^ cells could also be detected in p63^−^ cells, consistent with our immunofluorescence results (Fig. 2d-f and Supp. Fig. 3a, b). We characterized p63^+^Sox2^+^ and p63^−^Sox2^+^ cell populations in matured D3-organoids in detail. The p63^+^Sox2^+^ population could be further clustered into p63^+^Sox2^high^ subpopulation with high level of *Col17A1* gene expression and p63^+^Sox2^low^ subpopulation with high levels of *MKi67* and *Cdk1* cell cycle-related gene expression (Supp. Fig. 3c). However, these p63^+^Sox2^+^ cells were CK14^high^ cells, indicating that they represented the basal layer cells with stemness property (Col17A1^high^) and growth property (MKi67^high^), which consistent with our previous study [24]. In contrast, the p63^−^Sox2^+^ population were CK13^high^CK14^low/negative^ cells expressing high levels of differentiation-related genes, such as *Sprr1a*, suggesting that p63^−^Sox2^+^ cells were suprabasal cells. Moreover, the p63^−^Sox2^high^ cell population also displayed high levels of inflammation-related gene *DMKN*, *S100a8* and *S100a9* expressions, suggesting that the p63^−^Sox2^+^ cells could be involved in inflammatory stress response (Supp. Fig. 3c). Taken together, these results led us propose that the p63^+^Sox2^+^ cells presented in the basal layer of stratified “matured” D3-organoids were either the stem cells of origin responsible for maintaining cell stemness (p63^+^Sox2^+^ Col17A1^high^ cells) or proliferating cells of origin responsible for promoting cell proliferations (p63^+^Sox2^+^ MKi67^high^ cells). In contrast, the p63^−^Sox2^+^ cells presented in the suprabasal layers of stratified “matured” D3-organoids were the cell pool that represented differentiating cells of origin for meditating further cell differentiations (p63^−^Sox2^−^ cells and then keratinized layer) under normal conditions (Fig. 3a and Supp. Fig. 3). However, p63^−^Sox2^+^ might have a dual role responsible for rapid cell proliferations in tissue damage-repair responses under damage/stress conditions (Fig. 3a and Supp. Fig. 3c, e).

To test the possibilities, we suppressed p63 or Sox2 expression via RNA interference (RNAi) using lentivirus-mediated shRNA specifically targeting p63 or Sox2 in 2-D cultured D3 cells, respectively (Supp. Fig. 4a). Suppression of p63 expression led to cell differentiation and senescence, especially prohibitions of organoid formation in D3 cells. In contrast, suppression of Sox2 expression had less effects on cell growth in 2-D cultured D3 cells but also prohibited D3-organoid formation (Supp. Fig. 4b). These results indicated that p63 and Sox2 had essential roles for the stratified squamous epithelial formation and development, consistent with previous reported [39].

To avoid inhibitions of D3-organoid formation and development by p63 or Sox2 suppression, we grew D3-organoids to day 3.5, infected these organoids with p63-shRNA or Sox2-shRNA lentiviruses and then analyzed them at Day 10. Immunofluorescence analysis showed that p63 or Sox2 expression could be suppressed in D3-organoids by this strategy although suppression of p63 or Sox2 expression in D3-organoids were not as efficient as in 2-D cultured D3 cells (Fig. 3c, e). Nevertheless, suppression of p63 severely affected the basal layer cell maintenance in stratified “matured” D3-organoids, resulting in significantly decreased basal cells with undetectable levels of CK14 expression (Fig. 3d, e). Consistently, the suprabasal layer cells were also decreased without CK13 expression and the organoid sizes of p63-depleted organoids were significant smaller than those of controls (Fig. 3d, e, Supp. Fig. 4c).

Suppression of Sox2 in D3-organoids also significantly perturbed D3-organoid growth when compared with controls (Fig. 3b). However, unlike suppression of p63 expression resulting in inhibition of Sox2 expression, suppression of Sox2 in D3-organoids did not affect the expression of p63 in the basal cells in stratified “matured” D3-organoids (Fig. 3c, e). Although CK14^+^ cells and CK13^+^ cells were still detected in the basal layer and suprabasal layers, both layers in Sox2-depleted organoids formed disorganized structures with the morphologies of the suprabasal layers displaying more server aberrances than those of the basal layer (Fig. 3d, e). These results indicated that, while both pluripotent stemness factors, Sox2 and p63, were required for cell proliferation-differentiation homeostasis and stratified tissue maintenance of D3-organoids, Sox2 had its distinctive role(s) for regulation of the processes, especially in the suprabasal layers.

### Determination of p63^−^Sox2^+^ population as a critical cell pool in the suprabasal layers required for damage/stress responses in esophageal stratified epithelia in vitro and in vivo

To define the distinctive role(s) of p63^−^Sox2^+^ population in the suprabasal layers of D3-organoids, we set up an inflammatory stress response assay using the D3 rEO system. A detailed experimental flow chart is shown in Fig. 4a in that D3-organoids were divided into 4 groups: 1) control D3-organoids, which were grown for 10 days; 2-3) D3-organiods, which were grown for 10 days but at Day 7 or Day 5 treated with TNF-α and IL-13 for 3 days or 5 days to induce acute inflammatory stress responses; and 4) D3-organiods, which were grown for 10 days but at Day 5 treated with TNF-α and IL-13 for 3 days to induce acute inflammatory stress responses and then at Day 8 withdrawn the TNF-α and IL-13 treatment and then replaced into fresh medium for 2 more days allowing the acute inflammatory stress response recovery. Immunoblotting analysis indicated that treatment of D3-organoids with TNF-a and IL-13 resulted in increased expressions of acute inflammatory stress response proteins, S100a8 and S100a9 and the apoptosis-related proteins, Bax, cleaved-caspase3 and cleaved-PARP, confirming that TNF-α and IL-13 treatment induced inflammatory stress responses in D3-organoids (Fig. 4b).

**Fig 4.**
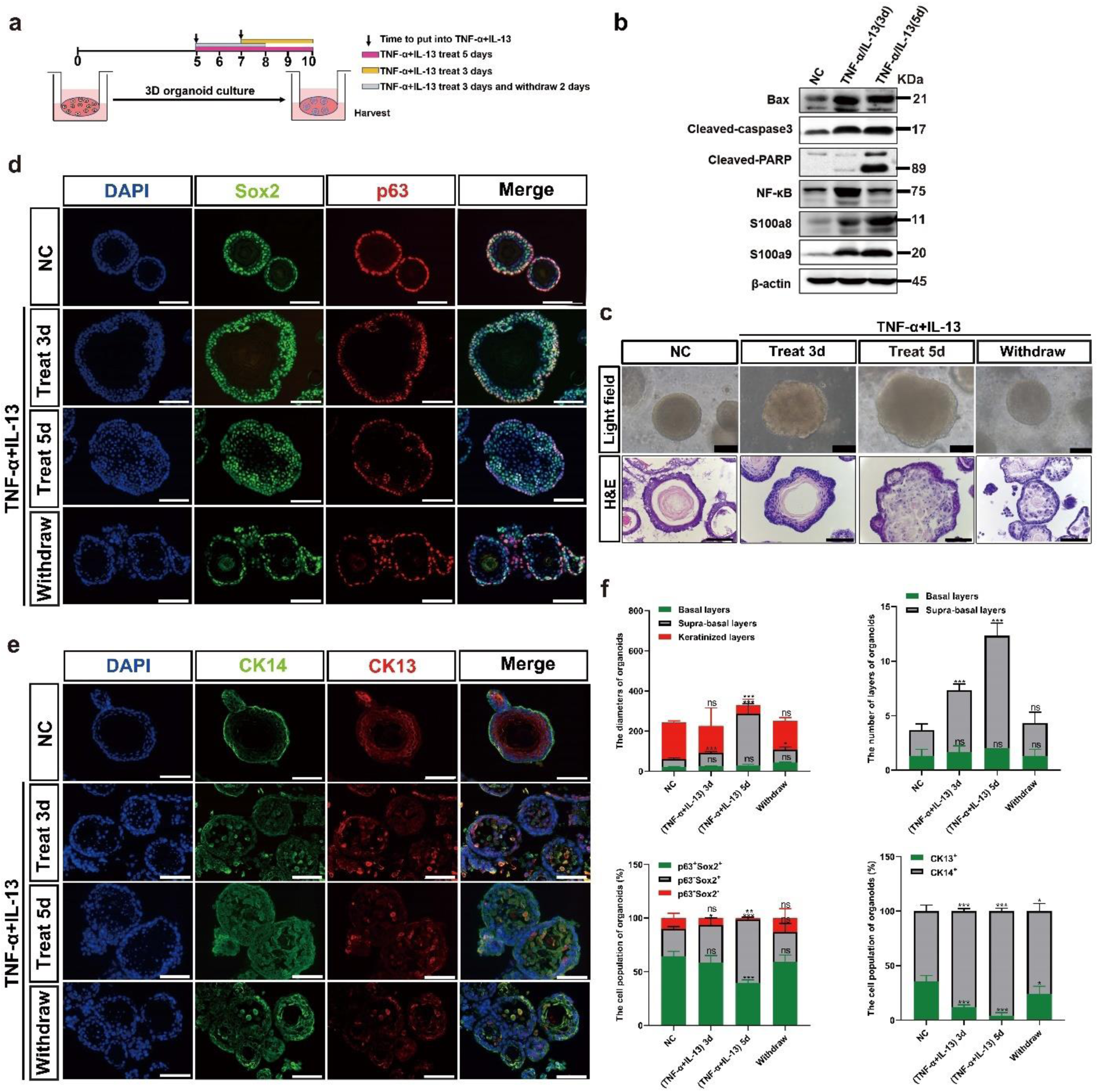
The p63^−^Sox2^+^ population responses damage/stress in esophageal stratified epithelia of D3-organoids. **a** Schematic illustration of the establishment of an inflammatory stress response assay using the rEO system. **b** Western blotting of inflammatory and apoptosis related proteins expression of D3-organoids treated with TNF-α (40 ng/mL) and IL-13 (20 ng/mL) at Day 10. **c** Representative brightfield, H&E staining images of D3-organoids treated with TNF-α (40 ng/mL) and IL-13 (20 ng/mL) at Day 10, Scale bar: 200 μm. **d** P63 (red) and Sox2 (green) coimmunostaining of D3-organoids treated with TNF-α (40 ng/mL) and IL-13 (20 ng/mL), Scale bar: 200 μm. All images are representative of three experiments. **e** CK13 (red) and CK14 (green) coimmunostaining of D3-organoids treated with TNF-α (40 ng/mL) and IL-13 (20 ng/mL) Scale bar: 200 μm. All images are representative of three experiments. **f** Quantifications of the diameters, layers number, cell number of p63^+^Sox2^+^, p63^−^Sox2^+^ and p63^−^Sox2^−^ populations, and cell number of CK13^+^ and CK14^+^ populations of D3-organoids treated with TNF-α (40 ng/mL) and IL-13 (20 ng/mL) (n=4, each n represents four random microscope fields, 400×). The data are presented as mean ± SD for percentage analysis, \**P*< 0.05, \*\**P* < 0.01, \*\*\**P*< 0.001, one-way ANOVA test on all pairwise combinations.

Consistently, phase-contrast microscopy revealed that, when compared to controls, treatments of D3-organoids with TNF-α and IL-13 for 3 or 5 days resulted in increasing organoid sizes and forming irregular nonsmoothed shapes of spheres (Fig. 4c). However, these D3-organoid abnormalities were quickly recovered when TNF-α and IL-13 treatment were withdrawn (Fig. 4c). Immunofluorescence analysis also showed that, when compared with controls, both p63^+^Sox2^+^ and p63^−^Sox2^+^ cell populations at the basal layer and suprabasal layers were increased in D3-organoids treated with TNF-α and IL-13 but p63^−^Sox2^+^ cell population proliferated more significantly than p63^+^Sox2^+^ cell population (Fig. 4d, f). The increasements of p63^−^Sox2^+^ population were not only accompanied with the significant increased the cell numbers and but also the layers of the suprabasal layers (Fig. 4d, f). Notably, increased p63^−^ Sox2^+^ cells in the suprabasal layers were CK14^+^ cells but not CK13^+^ cells (Fig. 4d, f), suggesting that the p63^−^Sox2^+^ cell population in stratified “matured” D3-organoids under the TNF-α and IL-13 treatment quickly converted from its differentiation-prone status to proliferation-prone status. Thus, the p63^−^Sox2^+^ cell population in the suprabasal layers of stratified “matured” D3-organoids could function as a cell pool that had a dual role required not only for mediating further cell differentiations but also for rapidly responding damage/stress assaults to maintain tissue homeostasis. Consistently, as TNF-α and IL-13 treatments were removal from D3-organoids, the increasements of p63^−^Sox2^+^ population were reduced, accompanying with reductions of the irregular D3-organoid shapes, sizes and the layers of stratified suprabasal layers (Fig. 4c-f).

To ascertain that p63^−^Sox2^+^ population functioned as a critical cell pool in the suprabasal layers responsible for damage/stress responses in esophageal stratified squamous epithelia, we depleted p63 or Sox2 expression from TNF-α and IL-13 treated D3-organoids by lentivirus-medicated RNAi as described above. Immunofluorescence analysis showed that treatment of TNF-α and IL-13 in Sox2-depletd organoids did not cause increased cell numbers and layers in the suprabasal layers when compared with controls, indicating that suppression of Sox2 expression, especially depletion of the p63^−^Sox2^+^ cell population in D3-organoids diminished inflammatory stress responses (Fig. 5a-d). In contrast, treatment of TNF-α and IL-13 in p63-depleted organoids resulted in increased cell numbers and layers in the suprabasal layers (Fig. 5a-d). However, unlike control D3-organoids, cells in Sox2-depleted D3-organoids treated with TNF-α and IL-13 were stained positively for CK14 and CK13 whereas cells in p63-depleted D3-organoids treated with TNF-α and IL-13 were only stained positively for CK13 (Fig. 5c, d), demonstrating that suppression of p63 or Sox2 expression also affected the stemness maintenance and cell proliferation-differentiation homeostasis in the basal cells.

**Fig 5.**
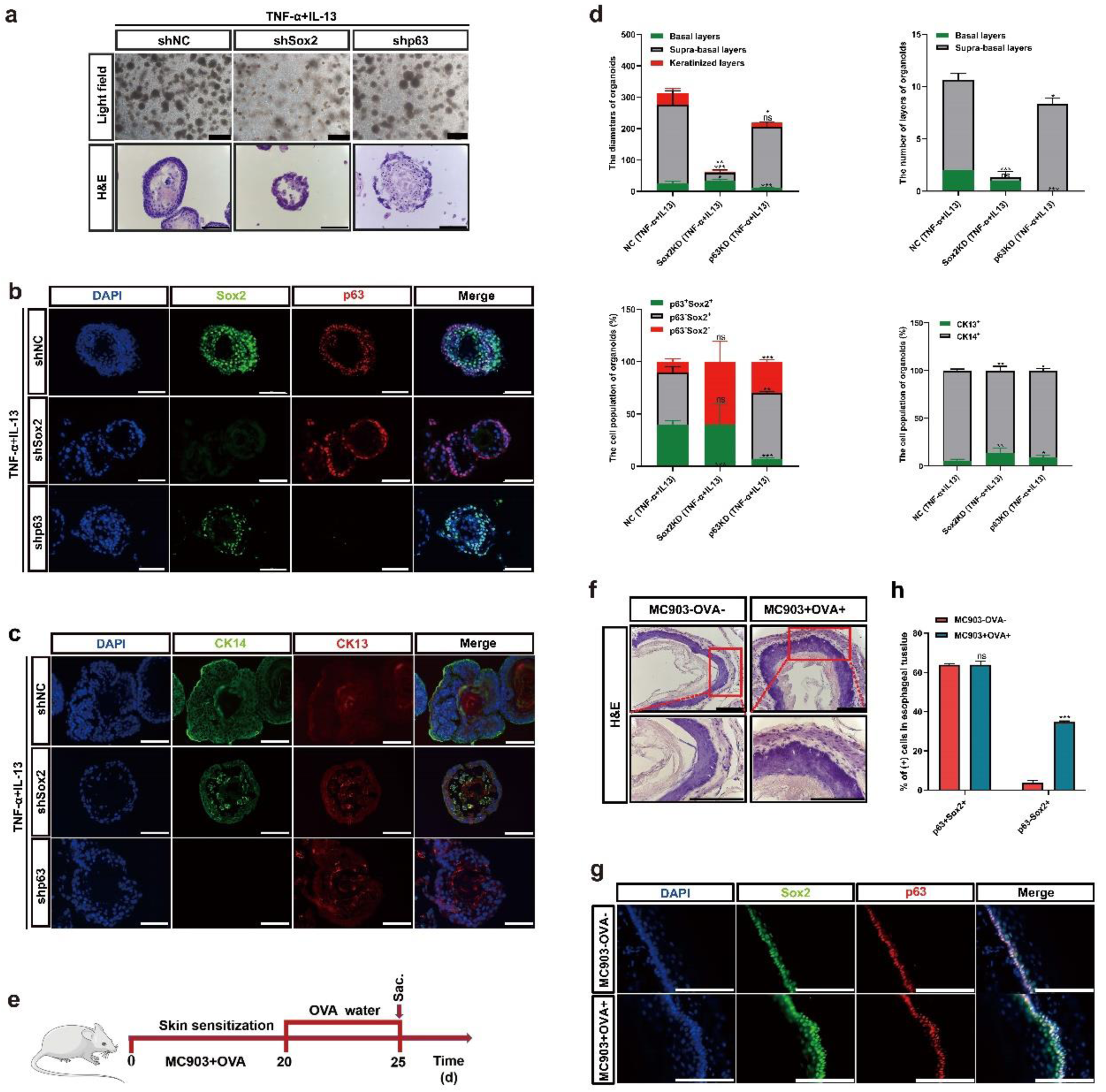
The p63^−^Sox2^+^ population as a critical cell pool in the suprabasal layers required for damage/stress responses in vitro and in vivo. **a** Representative brightfield, H&E staining images of TNF-α and IL-13 treated shp63 or shSox2 D3-organoids at Day 10, Scale bar: 200 μm. **b** P63 (red) and Sox2 (green) coimmunostaining of TNF-α and IL-13 treated shp63 or shSox2 D3-organoids, Scale bar: 200 μm. All images are representative of three experiments. **c** CK13 (red) and CK14 (green) coimmunostaining of TNF-α and IL-13 treated shp63 or shSox2 D3-organoids, Scale bar: 200 μm. All images are representative of three experiments. **d** Quantifications of the diameters, layers number, cell number of p63^+^Sox2^+^, p63^−^Sox2^+^ and p63^−^Sox2^−^ populations, and cell number of CK13^+^ and CK14^+^ populations of TNF-α and IL-13 treated shp63 or shSox2 D3-organoids (n=4, each n represents four random microscope fields, 400×). The data are presented as mean ± SD for percentage analysis, \**P*< 0.05, \*\**P* < 0.01, \*\*\**P*< 0.001, one-way ANOVA test on all pairwise combinations. **e** Schematic illustration of establishing rat inflammatory esophagitis model in vivo. **f** Representative H&E staining images of esophageal epithelial tissues, Scale bar, 100 μm, 200 μm. **g** P63 (red) and Sox2 (green) coimmunostaining of esophageal epithelial tissues, Scale bar: 200 μm. All images are representative of three experiments. **h** Quantifications of the percentage of p63^+^Sox2^+^ and p63^−^Sox2^+^ populations of esophageal epithelial tissues. (n=4, each n represents four random microscope fields, 400×). The data are presented as mean ± SD for percentage analysis, \**P*< 0.05, \*\**P* < 0.01, \*\*\**P*< 0.001, one-way ANOVA test on all pairwise combinations.

Previous study showed that esophageal stratified squamous epithelia in adult rats could be damaged by food allergens that induced inflammatory responses and caused inflammatory esophagitis [40]. Therefore, we employed food antigens, ovalbumin (OVA) and vitamin D analog MC903 to induce esophagitis in F344 adult rats. A detailed experimental flow chart is shown in Figure 5E. After exposure to the food allergens, F344 rats developed inflammatory esophagitis featuring increased thicknesses of esophageal stratified epithelia when compared with controls (Fig. 5f). Co-immunofluorescence staining using anti-p63 and Sox2 antibodies confirmed that p63^−^Sox2^+^ cell population was significantly increased in the suprabasal layers of esophageal stratified epithelia in MC903/OVA-treated rat when compared with those of untreated rats (Fig. 5g, h). These in vivo results further confirmed that the suprabasal layers of esophageal stratified squamous epithelia existed a unique cell pool, p63^−^Sox2^+^, that could transiently and rapidly switch from the “differentiation-prone fate” under normal conditions to the “proliferation-prone fate” under tissue damage/stress conditions for maintaining the stratified tissue homeostasis (Fig. 7d).

### Perturbation of tissue homeostasis maintenance in D3-organoids by Sox2 overexpression and p63^−^ Sox2^+^ cell population was undetectable in ESCC-organoids

Previous studies demonstrated that long-term damage/stress injuries at stratified esophageal epithelia could lead to develop ESCCs, and Sox2 was developmental lineage oncogene commonly overexpressing in human ESCCs [41]. As p63^−^Sox2^+^ cells were shown to have the dual role for maintaining tissue homeostasis and damage/stress responses in the suprabasal layers of esophageal stratified epithelia, we next asked whether overexpression of Sox2 in D3-organoids (SOE-D3-organoids) could lead to increase the p63^−^Sox2^+^ cell population that perturbed the tissue homeostasis maintenance and/or damage/stress responses in the suprabasal layers of stratified epithelia and whether the p63^−^Sox2^+^ cell population was involved in esophageal carcinogenesis. To this end, we transduced human HA-tagged Sox2 cDNA into 2-D cultures of D3 cells by lentivirus (Neo+)-mediated infection. After Neomycin selection, resistant cells were pooled and examined by immunoblotting analysis. As shown in Figure 6A, when compared with controls, SOE-D3 cells overexpressed ∼2 folds of exogenous HA-tagged Sox2, which also resulted in increased expressions of cyclin D1, ECT2 and CK14 and decreased expressions of KLF4 and CK13. In contrast, overexpression of Sox2 in D3 cells did not affect the p63 expression.

**Fig 6.**
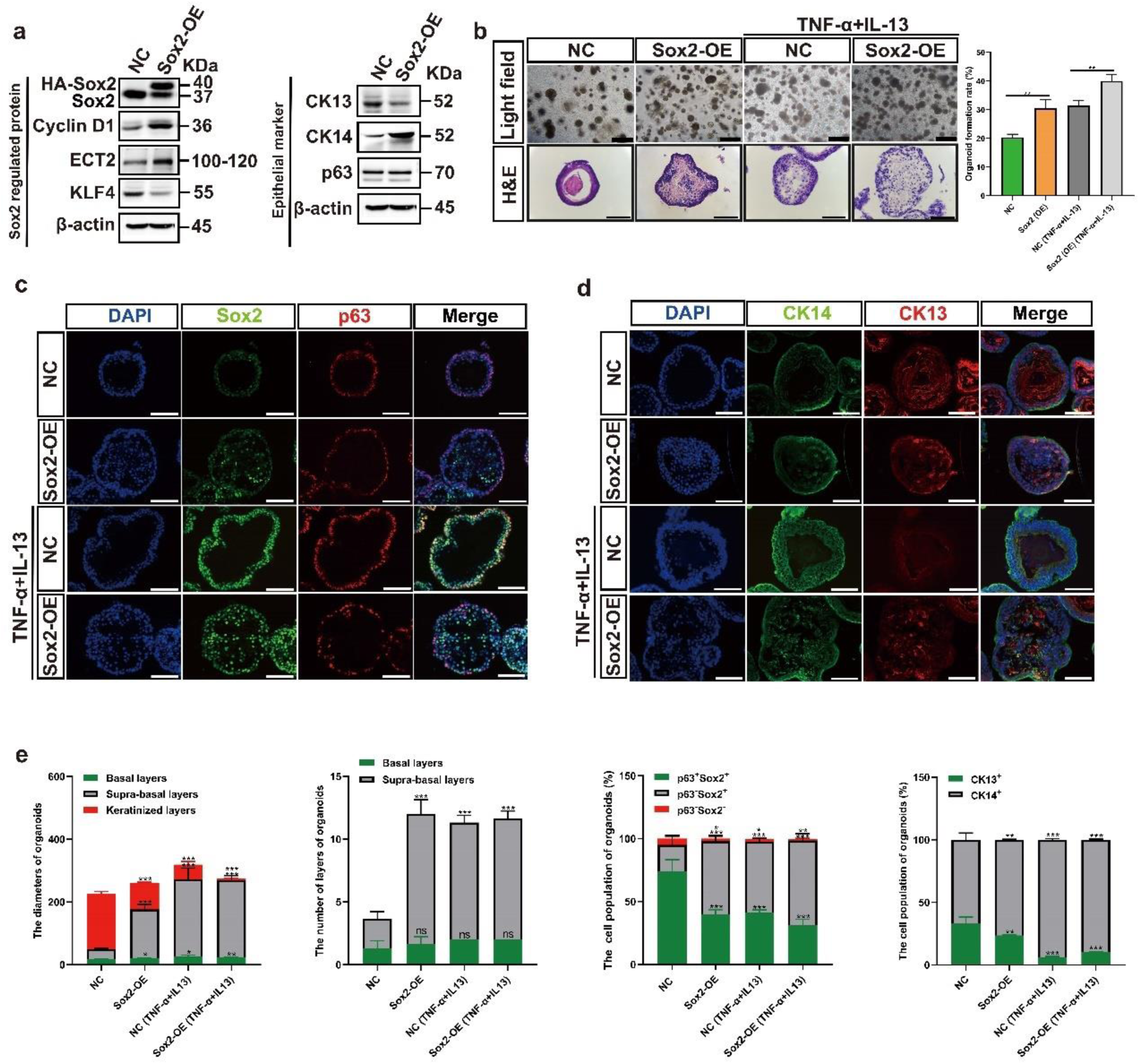
Overexpression of Sox2 perturbs tissue homeostasis maintenance in D3-organoids. **a** Western blotting of Sox2 regulated proteins and epithelial marks of SOE-D3 cells. **b** Representative brightfield, H&E staining images and quantification of OFR of SOE-D3-organoids and SOE-D3-organoids with TNF-α and IL-13 treatment at Day 10, Scale bar: 200 μm. **c** P63 (red) and Sox2 (green) coimmunostaining of SOE-D3-organoids and SOE-D3-organoids with TNF-α and IL-13 treatment, Scale bar: 200 μm. All images are representative of three experiments. **d** CK13 (red) and CK14 (green) coimmunostaining of SOE-D3-organoids and SOE-D3-organoids with TNF-α and IL-13 treatment, Scale bar: 200 μm. All images are representative of three experiments. **e** Quantifications of the diameters, layers number, cell number of p63^+^Sox2^+^, p63^−^Sox2^+^ and p63^−^Sox2^−^ populations, and cell number of CK13^+^ and CK14^+^ populations of SOE-D3-organoids and SOE-D3-organoids with TNF-α and IL-13 treatment (n=4, each n represents four random microscope fields, 400×). The data are presented as mean ± SD for percentage analysis, \**P*< 0.05, \*\**P* < 0.01, \*\*\**P*< 0.001, one-way ANOVA test on all pairwise combinations.

SOE-D3 cells were grown under the organoid culture condition and analyzed by phase-contrast microscopy, H&E-staining and immunofluorescence analyses. As shown in Figure 6B-6E, when compared with control D3-organoids, SOE-D3-organoids displayed enhanced OFR, formed irregular shapes and sizes and increased p63^−^Sox2^+^ cell population with CK14^+^ in the suprabasal layers, similar to those observed in D3-oragoids treated with TNF-α and IL-13. However, unlike D3-organoids treated with TNF-α and IL-13 resulting in significant decreases of CK13 cells in the suprabasal layers, SOE-D3-organoids still retained substantial CK13^+^ cells in the suprabasal layers, which were detected in control D3-organoids. These results indicated that overexpression of Sox2 in D3-organoids perturbed but did not completely block (or switch) the “differentiation-prone fate” of the p63^−^Sox2^+^ cells in the suprabasal layers. Consistently, p63^−^Sox2^−^ cells and the keratinized layer were still detected in SOE-D3-organoids (Fig. 6d, e).

Based on the results, one might expect that treatment of SOE-D3-organoids with TNF-α and IL-13 would induce severe inflammatory responses, resulting in greatly enhanced p63-Sox2+ cell population in the suprabasal layers of stratified “matured” epithelia. As shown in Fig. 6b-e, immunofluorescence analysis revealed that, when compared with D3-organoids, D3-organoids treated with TNF-α and IL-13 and SOE-D3-organoids, SOE-D3-organoids treated with TNF-α and IL-13 displayed severe hyperplasia determined by increasing p63^−^Sox2^+^ cells and decreasing p63^−^Sox2^−^ cells with disorganized tissue structures in the suprabasal layers. Consistently, increased p63^−^Sox2^+^ cells in the suprabasal layers were CK14^+^ cells but not CK13^+^ cells, and the keratinization process was also suppressed in SOE-D3-organoids treated with TNF-α and IL-13.

Given the fact that the tissue homeostasis maintenance and damage/stress responses were abrogated in cancers [42, 43], we finally explored whether the p63^+^Sox2^+^ and/or p63^−^Sox2^+^ cell populations would change in abnormal epithelia of ESCCs. Therefore, we generated ESCC organoids from a rat ESCC cell line, RESC-1, derived from a NMBzA-induced rat ESCC we previously described [44, 45]. As shown in Figure 7A, RESC-1-organoids were morphologically irregular that formed large, poorly differentiated cellular clusters with hyperchromatic nuclei and high nuclear-to-cytoplasmic ratio, consistent with advanced dysplasia/carcinoma phenotypes. Co-immunofluorescence analysis using anti-p63 and Sox2 antibodies in RESC-1 organoids demonstrated that the p63^−^Sox2^+^ cells were rarely detected in the abnormal epithelia of RESC-1-organoids. Instead, the p63^+^Sox2^+^ population occupied in almost the entire aberrant epithelial layers of RESC-1 organoids (Fig. 7a, b). Similar results were also observed in organoids derived from a human ESCC cell line, KYSE410 (Fig. 7c). These results were not unexpected because p63 and Sox2 functioned as cell lineage oncogenes commonly co-overexpressed in ESCCs and the highly “proliferating-prone” p63^+^Sox2^+^ population replaced the “differentiating-prone” p63^−^Sox2^+^ population, resulting in disruptions of the cell proliferation-differentiation and tissue homeostasis maintenance in abnormal ESCC epithelia (Fig. 7d).

**Fig 7.**
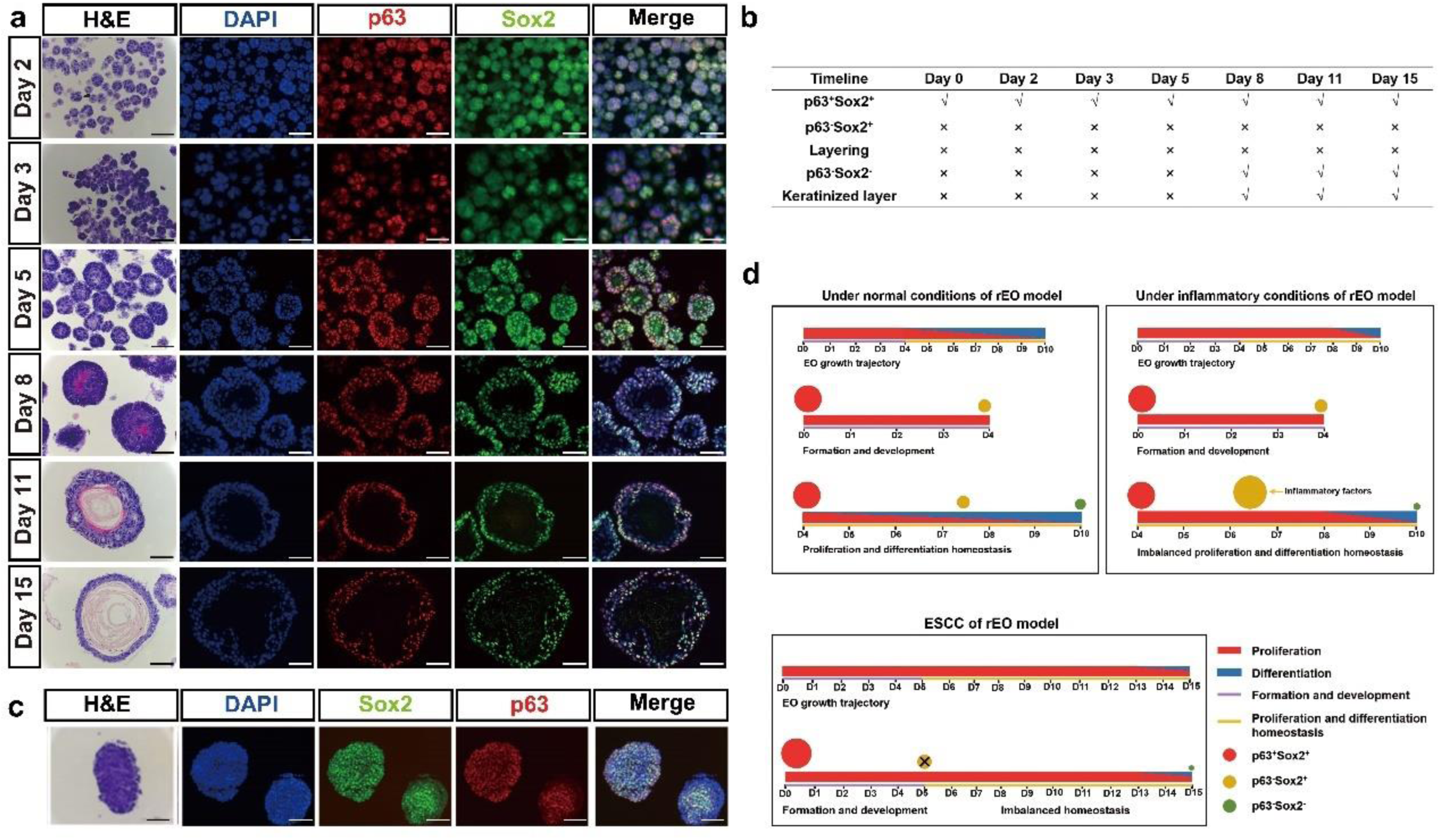
The p63^−^Sox2^+^ population is undetectable in ESCC-organoids. **a** Representative brightfield, H&E and immunostaining images of rat ESCC organoids derived from RESC-1 in formation trajectory, Scale bar: 200 μm. **b** The chart of features of rat ESCC-organoids by formation trajectory. **c** CK13 (red) and CK14 (green) coimmunostaining, P63 (red) and SOX2 (green) coimmunostaining of ESCC-organoids derived from human ESCC cell line KYSE410, Scale bar: 200 μm. All images are representative of three experiments. **d** Roles of p63^+^Sox2^+^, p63^−^Sox2^+^ and p63^−^Sox2^−^ populations in proliferation-differentiation homeostasis, self-renewal and maintenance, damage-repair/recovery, and carcinogenesis process of esophageal stratified epithelium.

## Discussion

In this study, we report the establishment of a stable in vitro rEO model system that enables us to investigate the cell proliferation-differentiation and tissue homeostasis maintenance in mammalian esophageal stratified “matured” epithelia, which have not been determined due to lack of available systems. The rEOs derived from single hTERT-immortalized “normal” rat esophageal keratinocyte D3 cells have a constant OFR and display cell expansion, epithelial layer formation, stratification and keratinization, similar to rat esophageal stratified squamous epithelia. By using the rEO system together with various methods including daily spatiotemporal snapshot imaging and the statistical analyses, we depict the landscapes of D3-organoid formation, development and maturation trajectories. Our results demonstrate that D3-organoids form from single p63^+^Sox2^+^CK14^+^ D3 cells to small asymmetrically p63^+^Sox2^+^CK14^+^ cell clumps; then p63^+^Sox2^+^CK14^+^ cell clumps grow into simple columnar epithelia with p63^+^Sox2^+^CK14^+^ and p63^−^Sox2^+^CK14^+^ cells presented; thereafter simple columnar epithelia develop into stratified multilayered epithelia with p63^+^Sox2^+^CK14^+^ cells presented in the basal layer and p63^−^Sox2^+^CK14^+^/p63^−^Sox2^+^CK13^+^ cells presented in the suprabasal layers; and finally mature into stratified multilayered epithelia with p63^+^Sox2^+^CK14^+^ cells presented in the basal layer, p63^−^ Sox2^+^CK13^+^/p63^−^Sox2^−^CK13^+^ cells presented in the suprabasal layers and the keratinization layer in the organoid centers. These results indicate that, under normal rEO culture conditions, the cell fate and commitment(s) for proliferation-differentiation occur at the beginning of organoid formation and cell lineages during organoid development and maturation are from p63^+^Sox2^+^CK14^+^ cells to p63^−^ Sox2^+^CK14^+^ cells, then to p63^−^Sox2^+^CK13^+^, p63^−^Sox2^−^CK13^+^ cells and the keratinization.

These results together with the scRNA-seq analysis, gene-level manipulations and damage-stress responses (Fig. 3–4 and Supp. Fig. 3) reveal that p63 and Sox2, two epithelial pluripotent determinants and developmental lineage oncogenes [11, 46–50], play crucial but distinctive roles in regulating cell proliferation-differentiation and tissue homeostasis of D3-organoid stratified epithelia. Notably, two cell populations, p63^+^Sox2^+^ in the basal layer and p63^−^Sox2^+^ in the suprabasal layers, which have not been previously defined, represent the cells of origin for proliferations and the cells of origin for differentiation in D3-organoid epithelia, especially stratified ‘matured” epithelia under normal culture conditions (Fig. 7d). Consistent with our results, previous studies demonstrated that p63 was only expressed in basal cells, which functioned as a determinant to control the commitment of early stem cells into basal cells progeny and the maintenance of basal cells in stratified squamous epithelia in vivo [15, 51, 52]. Moreover, p63 severed as a crucial “switch” that can cooperate with Sox2, another stratified squamous epithelial determinant, required for the basal layer to suprabasal layers cell lineage specifications [8, 53–56]. Thus, the p63^+^Sox2^+^ population presented at the basal layer is required for esophageal epithelial stemness maintenance and proliferations, which allow esophageal epithelia to constantly self-renew and to stratify into suprabasal layers.

In contrast, the p63^−^Sox2^+^ cells are only detected in the suprabasal layers of stratified “matured” epithelia of D3-organoids. Although Sox2^+^ cells were reported in the suprabasal layers of esophageal stratified epithelia by several studies previously, their role(s) had not been determined [8, 9]. However, Liu et al, used mouse genetic model to conditionally overexpress Sox2 in esophagus that led to epithelial hyperplasia with increase of cell proliferation and inhibition of cell differentiation in vivo [50]. Recently, the p63^−^Sox2^+^ cells were also identified in trachea pseudostratified columnar ciliated epithelia [57]. By landscape depiction of D3-organoid formation, development and maturation together with suppression or overexpression of Sox2 in D3-organoids, we demonstrate that the p63^−^Sox2^+^ cells presented in the suprabasal layers have a dual role to maintain cell proliferation-differentiation and tissue homeostasis of stratified epithelia. Under normal conditions, p63^−^Sox2^+^ cells are required for stratified epithelial further differentiations, but, under damage/stress conditions, the p63^−^Sox2^+^ cells switched its differentiation-prone fate to its proliferation-prone fate to rapidly facilitate stress/damage responses. Given the fact that esophagus in the gastrointestinal tract is frequently associated with chronic stress and injury during life, such as inflammatory and mechano-damages, causing frequently injury-repair processes [53], our studies explain how cells in the suprabasal layers are involved in controlling cell proliferation-differentiation and maintaining tissue homeostasis in esophageal stratified epithelia under normal and damage-stress conditions.

Hence, we hypothesize that the p63^+^Sox2^+^ population in the basal layer is essential for preserving esophageal epithelial stemness and promoting cell proliferations to suprabasal layers whereas the p63^−^ Sox2^+^ population in the suprabasal layers derived from the p63^+^Sox2^+^ population is responsible for mediating further cell differentiations and monitoring endogenous/exogenous strass-damage responses. Consistent with the hypothesis, we also show that only the p63^+^Sox2^+^ cells but not p63^−^Sox2^+^ cells are detected in ESCC organoids since cancer cells in abnormal ESCC epithelia perturb the cell proliferation-differentiation and tissue homeostasis maintenance and disrupt rapid damage-stress responses for advancing cell proliferations and tumorigenesis (Fig. 7d). In the future, our study should be verified by more sophisticated time lapse microscopy, such as the light-sheet microscopy even though the long-term live imaging of an organoid remains very challenging [58]. Escalations of the rEO system by genetic modifications, e.g., labeling D3 cells by knocking in GFP in lineage specific gene loci (p63, Sox2, CK13 or CK14) using CRISPR [59] and/or applications of advanced microscopic imaging technologies together with artificial intelligent (AI) deep learning [60] would help to direct visualize and trace cell lineages and determine cells of origin for stem maintenance, proliferations and differentiations in D3-organoids at the single cell level. Moreover, large scale scRNA-seq analyses of D3-organoids at the different stages with gene level manipulations would precisely determine what factors are crucial for determinations of proliferation-differentiation cell lineages switches of p63^+^Sox2^+^ cells to p63^−^Sox2^+^ cells under normal conditions and differentiation-prone p63^−^Sox2^+^ cells to proliferation-prone p63^−^Sox2^+^ cells under strass/damage conditions. The studies could also reveal when and how these factors control the cell proliferation-differentiation and tissue homeostasis maintenance in esophageal stratified epithelia at the molecular levels.

In summary, we develop a stable in rEO model system to investigate the cell proliferation-differentiation and tissue homeostasis maintenance of esophageal stratified epithelia, demonstrating that mammalian esophageal stratified tissue homeostasis is maintained distinctively by the epithelial pluripotent p63^+^Sox2^+^ and p63^−^Sox2^+^ cell populations in the basal layer and suprabasal layers. Our study has paved a way for the beginning of uncovering the cells of origin and cell lineages in mammalian esophageal stratified squamous epithelia.

## Acknowledgments

During this study (in 2019), Prof. Shih-Hsin Lu passed away. We all miss him.

## Abbreviation

rEO: Rat esophageal organoid
scRNA-seq: Single-cell RNA sequencing
ESCC: Esophageal squamous cell carcinoma
RES: Respiratory-esophageal separation
SRY: Sex-determining region Y
HDs: Hemidesmosomes
mEOs: Mammalian esophageal organoids
PSCs: Pluripotent stem cells
EoE: Eosinophilic esophagitis
SOE: Overexpression of Sox2
hTERT: Human telomere reverse transcriptase
RNE-D3: Immortalized rat normal esophageal epithelial D3 cell line
RESCs: Rat esophageal squamous carcinoma cell lines
FBS: Fetal bovine serum
OFR: Organoid formation rate
PCs: Primary rat esophageal squamous keratinocyte cells

## Statements and Declarations

### Author Contributions

All authors contributed to the study conception and design. Material preparation, data collection and analysis were performed by Xiaohong Yu and Hui Yuan. The first draft of the manuscript was written by Xiying Yu and Wei Jiang. All authors commented on previous versions of the manuscript. All authors read and approved the final manuscript.”

### Data Availability

Data will be made available upon reasonable request.

### Ethical approval

All experiments involving animals were complied with the standards approved by ethical committee of National Cancer Center/National Clinical Research Center for Cancer/Cancer Hospital, Chinese Academy of Medical Sciences.

---

**Supp. Fig 1.**
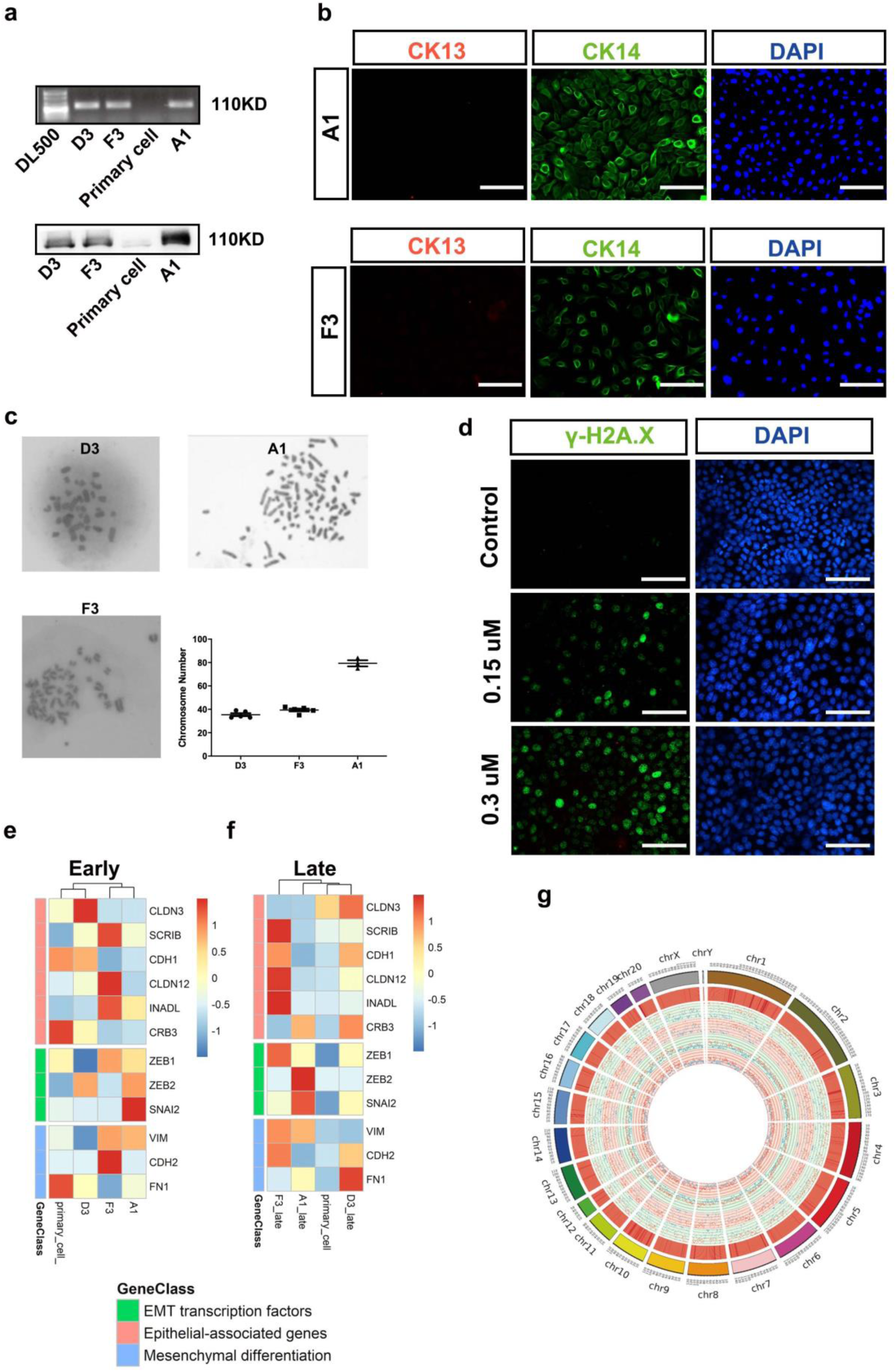
Generation of hTERT immortalized normal rat esophageal keratinocyte cell lines. **a** Detection of hTERT in RNE cell lines D3, F3, and A1. Electrophoresis of PCR products (top), Western blotting (bottom). **b** CK13 (red) and CK14 (green) coimmunostaining of F3 and A1 cell lines, Scale bar: 100 μm. All images are representative of three experiments. **c** Representative images of karyotype analysis and the statistics of D3, F3, and A1 cell lines. D3 (top left, 40×), A1 (top right, 100×) and F3 (bottom left, 40×). **d** Immunofluorescence staining of γH2A.x in D3 cell lines with adriamycin treatment after 12 hours, Scale bar: 100 μm. **e** Heatmap of epithelial-related gene expression in early passage of D3, F3 and A1 cell lines (passage 15-25) with primary rat esophageal epithelial cells as control. **f** Heatmap of epithelial-related gene expression in lately passage of D3, F3 and A1 cell lines (passage > 50) with primary rat esophageal epithelial cells as control. **g** Circle plot of whole genomic sequencing of D3 cell line.

**Supp. Fig 2.**
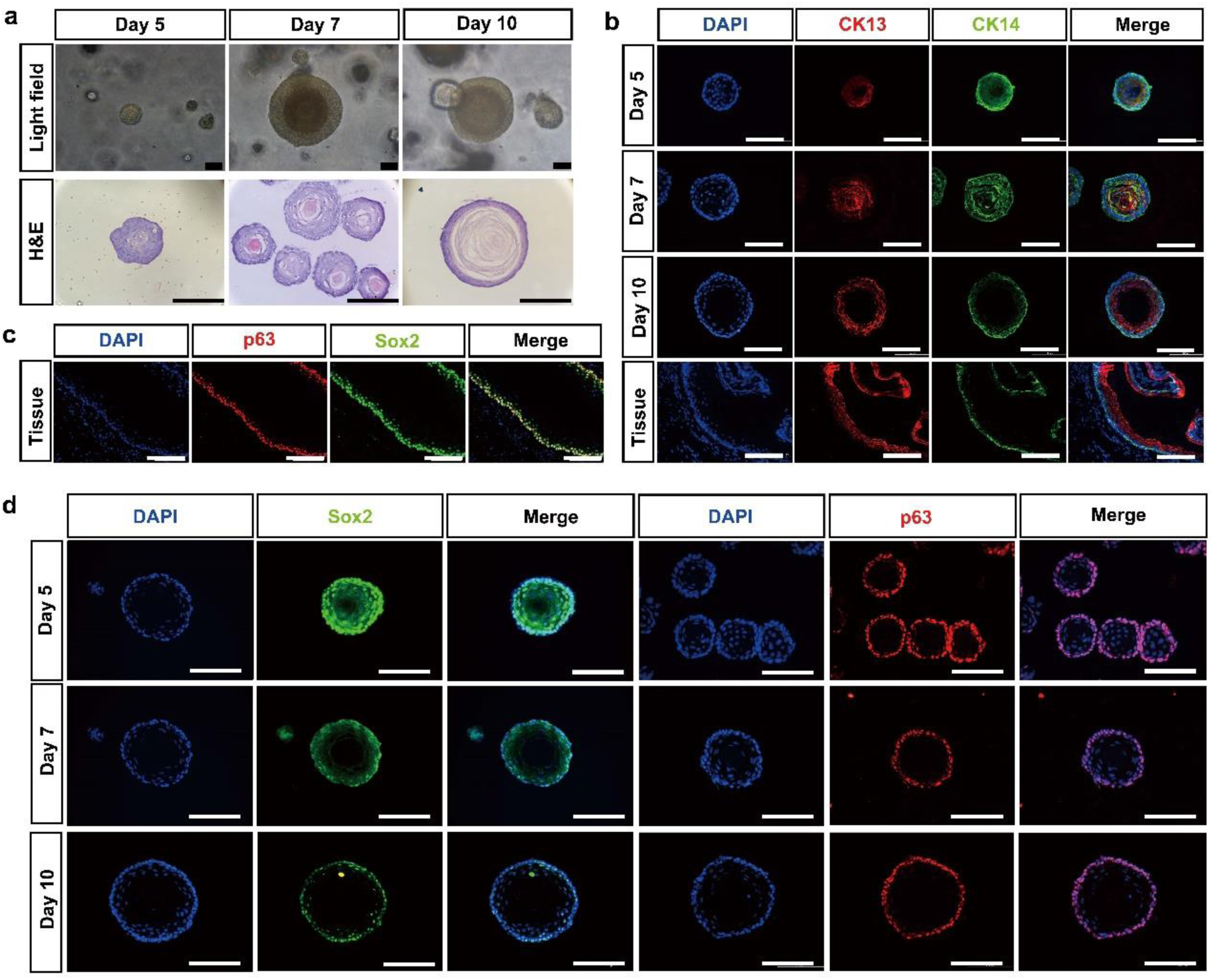
Characterization of rat esophagus and esophageal organoids derived from primary rat esophageal basal keratinocyte cells. **a** Representative brightfield, H&E staining images of organoids derived from primary rat esophageal basal keratinocyte cells in formation trajectory, Scale bar: 200 μm. **b** CK13 (red) and CK14 (green) coimmunostaining in rat esophagus and esophageal organoids derived from primary rat esophageal basal keratinocyte cells in formation trajectory, Scale bar: 200 μm. **c** P63 (red) and Sox2 (green) coimmunostaining of rat esophagus, Scale bar: 200 μm. **d** P63 (red) and Sox2 (green) coimmunostaining in rat esophagus and esophageal organoids derived from primary rat esophageal basal keratinocyte cells in formation trajectory, Scale bar: 200 μm.

**Supp. Fig 3.**
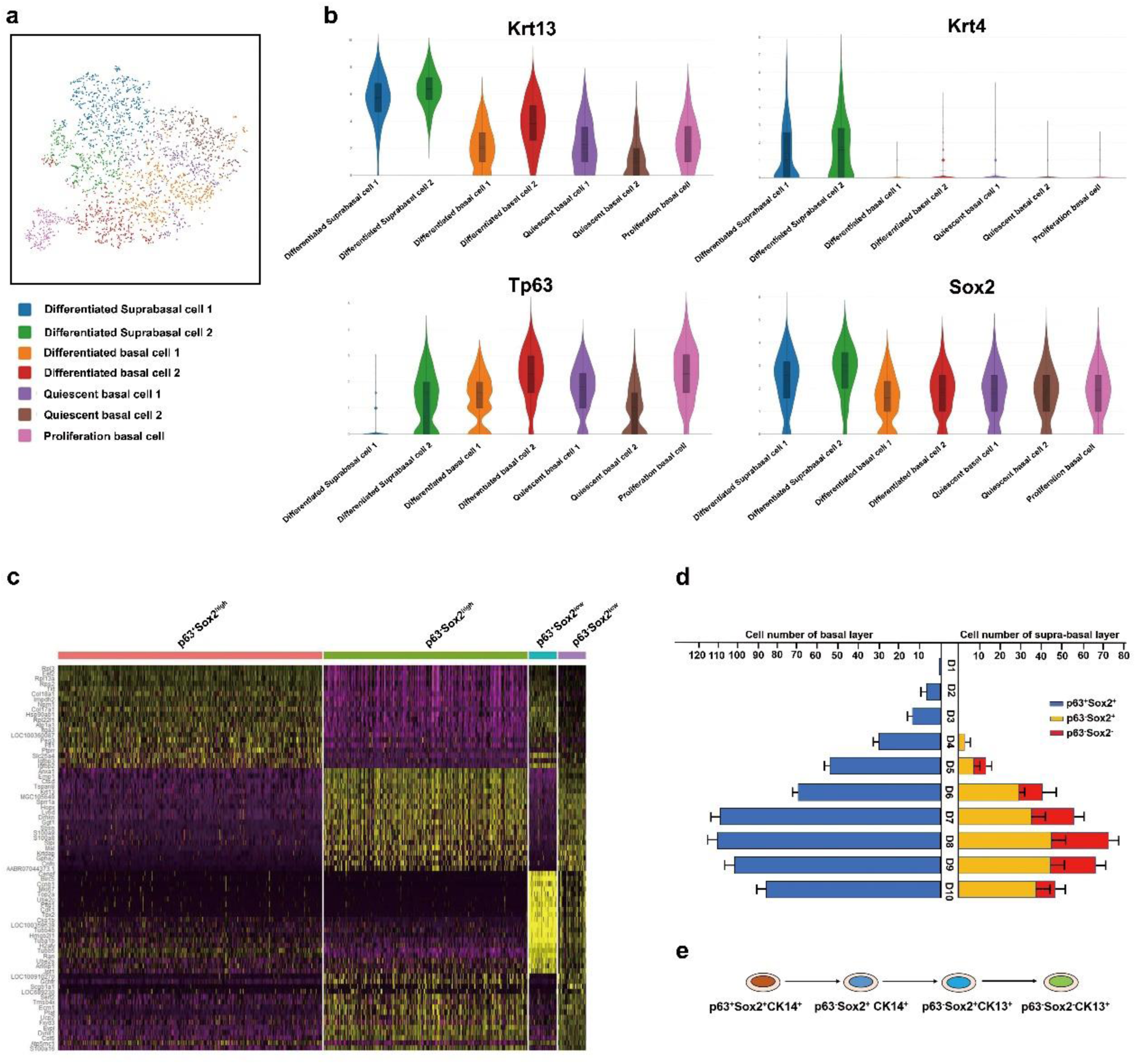
scRNA-seq analysis of rat esophageal organoids derived from D3 cells. **a** t-SNE plot displaying the scRNA-seq data. Different colors indicate distinct cell subpopulations. **b** Violin plots of Krt13, Krt4, Sox2 and p63 gene expression among the seven different subpopulations. The y-axis represents the expression levels of the genes, and the x-axis represents the different subpopulations. **c** Heatmap showing the expression of selected genes from the four clusters corresponding to the subpopulations based on p63 and Sox2 gene expression. **d** Schematic illustration of the cell number of p63^+^Sox2^+^, p63^−^Sox2^+^ and p63^−^Sox2^−^ populations in the trajectory of esophageal organoids (n = 4, each n represents four random microscope fields, 400×). The data are presented as mean ± SD for percentage analysis. **e** The differentiation process of p63 and Sox2 dependent lineage of D3-organoids, \**P*< 0.05, \*\**P* < 0.01, \*\*\**P*< 0.001, one-way ANOVA test on all pairwise combinations.

**Supp. Fig 4.**
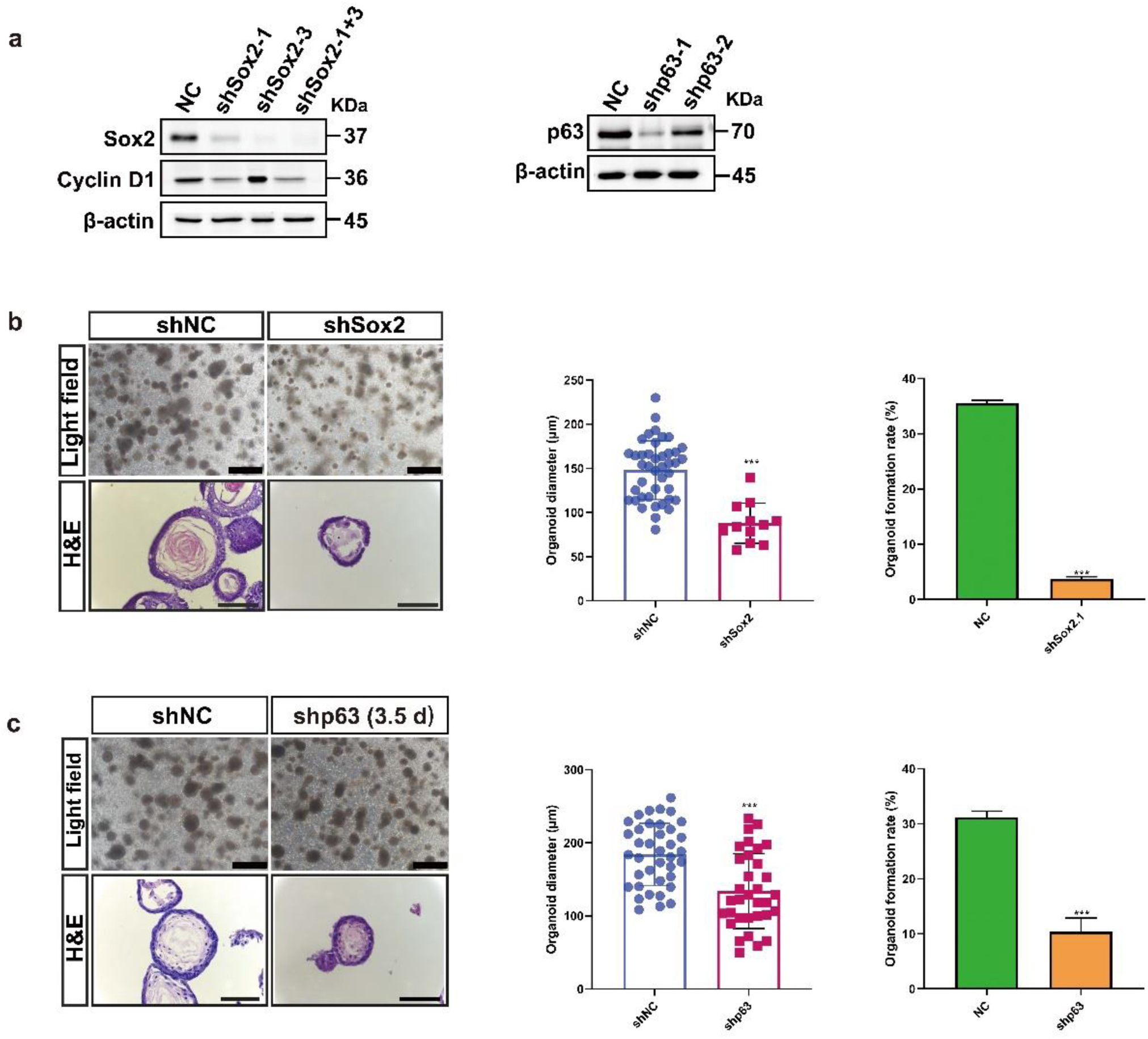
Knockdown of p63 and Sox2 in D3 cells in 2-D and in D3-organiods in 3-D. **a** Western blotting verification of Knockdown of p63 and Sox2 by lentivirus in D3 cells in 2-D. **b** Representative brightfield and H&E staining images. Quantification of diameters and OFR of −D3-organoids derived from Sox2 knockdown of D3 cells in 2-D, Scale bar: 200 μm. **c** Representative brightfield and H&E staining images. Quantification of diameters and OFR of D3-organoids that knockdown p63 at Day 3.5, Scale bar: 200 μm.

